# Orchards and paddy differentially impact rock outcrop amphibians: Insights from community- and species-level responses

**DOI:** 10.1101/2023.10.03.560737

**Authors:** Vijayan Jithin, Manali Rane, Aparna Watve, Rohit Naniwadekar

**Author notes:** Correspondence: Rohit Naniwadekar, Nature Conservation Foundation, Mysore, Karnataka, India.; ORCID ID: https://orcid.org/0000-0002-9188-6083 Vijayan Jithin, Nature Conservation Foundation, Mysore, Karnataka, India.; ORCID ID: https://orcid.org/0000-0002-9663-1661.

## Abstract

With agricultural demands increasing globally, determining the nature of impacts of different forms of agriculture on biodiversity, especially for threatened vertebrates and habitats, is critical to inform land management. We determined the impacts of converting rock outcrops (a habitat more threatened than rainforests) to orchards and paddy on anurans in the Western Ghats biodiversity hotspot. We sampled 50 belt transects four times across four sites during the rainy season and recorded information on amphibians and their microhabitats. We determined community-level responses using Hill numbers, beta-diversity measures, and non-metric multidimensional scaling, and species-level responses using joint species distribution modelling. Converting rock outcrops to paddy and orchards significantly altered microhabitat availability. Conversion to paddy mostly had community-level impacts, i.e., lowered species richness and more nested communities, whereas conversion to orchards mostly had species-level impacts, i.e., lowered species occurrence, highlighting the differential impacts of different forms of agriculture on amphibians and the need to determine impacts of land-use change on communities and species concurrently. We show that large rock pools are critical microhabitats for anurans as they serve as a refuge and protect anurans from desiccation during dry spells, which may be prolonged by climate change. Since rock outcrop habitats in low elevations are rapidly being converted to orchards, efforts are needed to conserve them in partnership with local communities, the custodians of these habitats. Our findings demonstrate that different forms of agriculture can have divergent impacts on biodiversity, and determining their impacts may require assessments at multiple scales, from species to communities.

## HIGHLIGHTS

1. Orchard and paddy conversions affect anurans and their microhabitats on rock outcrops
2. Conversion to paddy has community-level impacts, i.e., reduced diversity
3. Conversion to orchards has species-level impacts, i.e., lowered species occurrence
4. Rock pools, an important habitat for outcrop anurans, must be preserved or created
5. There is a need to study species- and community-level impacts simultaneously

## 1. INTRODUCTION

Conversion of natural habitats to agriculture is one of the primary drivers of biodiversity declines in the tropics (Newbold et al., 2015). Agricultural expansion is an expected outcome of increasing human consumption, associated with increasing population size and economic development (Laurance et al., 2014). Determining impacts of different forms of agriculture on biodiversity is vital for developing sustainable land-use management practices in the tropics, where agricultural demands are increasing and land-use changes are driving biodiversity declines (DeFries and Rosenzweig, 2010; Newbold et al., 2020). Fine-scale agroecological understanding is required to inform land-use management. This is critical for agroforestry plantations, as they are often recommended as alternative land-use production systems and habitats for biodiversity outside protected areas in the tropics (Bhagwat et al., 2008).

Different forms of agriculture, varying in the intensity of habitat modification, are often grouped while determining their impacts on biodiversity (Gonthier et al., 2014; Newbold et al., 2015). This may mask the contrasting effects of different forms of agriculture. Agriculture activities varying in intensity may have differing impacts, ranging from negatively impacting species abundances and filtering narrowly distributed specialist species to causing biotic homogenisation in extreme cases (McKinney and Lockwood, 1999; Solar et al., 2015). While certain land-use conversions may affect the abundance of particular species without necessarily affecting species richness, others act as filters of specialist species (Cingolani et al., 2007; Jesse et al., 2018). Therefore, it is critical to determine the specific impacts of different forms of agriculture on biodiversity to inform land management policy and practise better.

There is generally a bias towards the usage of community-level metrics (e.g., richness, composition) rather than investigating species-level responses when assessing the impacts of ecological perturbations (Kéfi et al., 2019). Changes in community composition can be due to turnover (species replacement), nestedness (species loss/gain), and variation in species abundance (Baselga, 2013; Baselga and Orme, 2012). Therefore, it is crucial to assess species- and community-level responses simultaneously. While community-level assessments provide information regarding overall impacts on perturbations on biodiversity, species-level assessments enable the determination of differences in species responses to perturbations. This aspect is valuable for developing species-specific conservation strategies. Very few studies have simultaneously examined species- and community-level responses to human activities (Asad et al., 2021; Fulgence et al., 2022).

Among vertebrates, amphibians are the most threatened and the loss of natural habitat is a primary threat to amphibian diversity, affecting 60% of all amphibians (Vié et al., 2009) and land-use change is one of the biggest threats to amphibian populations, particularly in south Asia (Cordier et al., 2021; Hof et al., 2011). Amphibians with narrow geographic distributions are vulnerable to habitat modification (Nowakowski et al., 2017). Ranges of almost 25% of amphibians remain outside protected areas (Nori et al., 2015). While in private lands, land-use change and associated habitat modification are inevitable, it is critical to determine the factors that can facilitate amphibian persistence and diversity in modified landscapes. Accordingly, understanding the species and community-level responses of amphibians to agricultural land-use is critical in the unprotected, modified landscapes. While studies have examined the impact of agricultural land-use changes on amphibians in forested habitats (Cordier et al., 2021), information from non-forested biomes, such as open natural ecosystems, is lacking.

Non-forested open ecosystems are among the most underappreciated and threatened ecosystems that support unique biodiversity and provide livelihoods for millions of pastoralists (Bond, 2019). Loss of open ecosystem habitats supersedes that of tropical rainforest biomes (Parr et al., 2014; Veldman et al., 2015). Often considered wastelands, these open ecosystems harbour rich diversity, which can be vulnerable to conversion to different land uses. The rock outcrops or lateritic plateaus of the northern Western Ghats are best examples for such threatened open ecosystems, as they harbour endemic biodiversity but are classified as ‘wastelands’ in official records (Government of India, 2019). In such seemingly bare landscapes, microhabitats can play a critical role in facilitating the persistence of biodiversity. However, our understanding of the role of different microhabitats in influencing species persistence is relatively poor.

Microhabitats, such as loose rocks, that shelter animals from extreme heat and rain play a critical role in harbouring diverse animal taxa (Jithin et al., 2023). Rock outcrops on the west coast of India are bare and dry in the summer but transform into an aquatic habitat with the onset of the monsoon, providing critical microhabitats for amphibians. A diverse array of habitats, including rock pools and flowing surface water, offer unique microhabitats for amphibians, including multiple endemic species, to thrive (Thorpe et al., 2018). While the ecology of these outcrops is poorly understood, rapid agricultural land-use changes are transforming such landscapes (Kulkarni et al., 2022; Madhusudan and Vanak, 2022). These plateaus have traditionally been converted to paddy, and mango and cashew plantations recently, differentially impacting native biodiversity (Bhattacharyya et al., 2019; Jithin et al., 2023). Conversion of these rock outcrops to paddy and agroforestry plantations offers contrasting examples of disturbances, one where the open ecosystem is converted to tree-based agriculture, and paddy wherein the surface is inundated with water for a prolonged duration. Given that these unique habitats are unprotected, privately owned, and vulnerable to land-use change, it is critical to determine the determinants of diversity and species persistence to inform land management policy. Amphibians on these rock outcrops are an excellent system to understand the species- and community-level responses to different forms of agriculture. While previous studies have examined the impacts of agroforestry plantations on forest-dwelling amphibians (Komanduri et al., 2023; Sankararaman et al., 2021), such information is lacking for open ecosystems.

Given this background, we investigated the impacts of land-use change on amphibians of rock outcrops in the northern Western Ghats. Across the land-use types, natural rock outcrops (henceforth, plateaus), agroforestry plantations (henceforth, orchards) and paddy, we compared: 1) the availability of microhabitats, 2) community-level responses (community composition, alpha- and beta-diversity, and overall abundance), and 3) species-level responses (probability of occurrence) of amphibians. Given the differential nature of conversion, we expected that 1) microhabitat availability will differ significantly among habitats, 2) conversion to paddy will result in lower diversity and more nestedness among communities due to species filtering and biotic homogenisation, and 3) conversion to orchards will reduce the availability of microhabitats, thereby negatively influencing frog species occurrence. By understanding the relationship between land-use change, microhabitat availability, and frog occurrence and diversity, we aimed to provide insights into maintaining or improving amphibian biodiversity in the modified landscapes (Smith et al., 2020).

## 2. MATERIALS AND METHODS

### 2.1. Study Area

We conducted the study in the low-elevation lateritic plateaus of the Ratnagiri region of Maharashtra State (16°31’–16°48’N; 73°19–73°29’E; Fig. S1), which forms the northern part of the Western Ghats, one of the eight ‘hottest’ biodiversity hotspots (Myers et al., 2000). These plateaus are privately owned or are under the government revenue department. The elevation of sampled plateaus was in the range 24–197 m asl. The area experiences a tropical climate (average rainfall: 3,313 mm; temperature range 23–33^°^C). The rainfall is restricted to the southwest monsoons (June–September). Heavy rains transform the dry habitat into a green carpet of herbaceous vegetation, a significant proportion of which is endemic (Watve, 2013) (Fig. S1). Thus, the fauna on the plateaus are exposed to hot and dry summers and water-logged monsoons (Watve, 2013). The microhabitats on these rock outcrops, such as loose rocks, pools, streams, and flush vegetation, provide refuge to multiple fauna (Jithin et al., 2023; Thorpe and Watve, 2015).

The study area is a mosaic of plateaus, orchards, paddy fields, and villages. Paddy is cultivated in the landscape traditionally by using the natural depressions with existing soil or by dumping soil on the plateau and lining it with boulders to prevent soil run-off. Paddy cultivation remains a significant agricultural land use in the landscape. The orchards on the plateaus generally consist of mango and cashew trees planted in large pits built by blasting the plateau, filling soil, and lining it with rocks (Bhattacharyya et al., 2019). The mango variety grown locally is known as ‘Ratnagiri Alphonso’ and is registered with a Geographical Indication tag. This variety, when grown on plateaus, is thought to fruit at desired times, well before the onset of monsoon, and is supposedly sweeter, thereby fetching a higher price (Bhattacharyya et al., 2019; Ganeshmurthy et al., 2018). Therefore, orchards are rapidly expanding on the plateau and are an important source of income (Bhattacharyya et al., 2019). Our study sites spanned these orchards, paddy fields, and unmodified plateaus (reference sites).

### 2.2. Sampling

Between June and September 2022, a period that coincides with the monsoon season, we sampled four plateaus: Devi Hasol, Devache Gothane, Gaonkhadi, and Bakale to capture the spatial variability (Fig. S1). We conducted nighttime belt (100 × 6 m^2^) transect surveys for amphibians (Scott et al., 1994) following Thorpe et al. (2018). We ensured no habitat differences or human disturbances (e.g., roads) within a transect. Fifty transects were laid out across four unique plateaus and three land-use types (20 on plateaus, and 15 each in paddy and orchards) (Fig. S1; Table S1). All transects were surveyed monthly (four temporal replicates), except the five orchard transects of Devache Gothane, where two temporal replicates could not be carried out due to political protests against a proposed refinery.

One observer (VJ) conducted the searches between 1730 and 0000 hr, usually in a clear climate, barring occasional rain incidences. At each 20 m point during each temporal replicate, in a 3-m radius circular subplot, the observer recorded the following microhabitat variables: rock- and shallow-pool volumes (cm^3^), maximum paddy depth (cm), stream cross-sectional volume (cm^3^), woody vegetation (%), flush vegetation (%), grass cover (%), and surface water presence. We calculated pool volume by multiplying the pool depth with the length (longest dimension) and width (second longest dimension) of the pool. We calculated stream cross-sectional volume within the subplot by multiplying the maximum depth and subplot diameter. The percentages of woody vegetation, flush vegetation, and grass cover were visually estimated. The observer recorded all visible amphibians.

### 2.3. Functional Trait Data

We compiled trait information for nine species, which occurred in at least 5% of the transects surveys using original species descriptions from published literature, if available. For species without published trait information, measurements from specimens at the Bombay Natural History Society (BNHS) Museum, India, were obtained (Saunak Pal, personal communication) (Table S2). Only male specimens were measured since female specimens were not available for all species. The traits included snout-vent length (SVL; body size), relative hindlimb length ([Femur+Shank length]/SVL), eye position (interorbital distance/head width), relative eye size (eye diameter/SVL), head shape (head length/head width), and degree of webbing (Garg and Biju, 2017). The latter two were excluded from the final analyses since they were strongly correlated (*r* > 0.7) with the other variables. Body size, eye size and position influence the foraging habit, limb length influences dispersal ability, and webbing influences the degree of dependency on aquatic habitats (Table S2).

### 2.4. Analyses

We performed all the analyses in R (v. 4.3.0) (R Core Team, 2023). To understand how the microhabitat characteristics differed between the land-use classes, we carried out unconstrained principal coordinate analysis (PCoA) on the Gower distances among transects based on the habitat characteristic data, using *cmdscale* function of the default package ‘stats’, and *vegdist* and *envfit* functions of the package ‘vegan’ (Oksanen et al., 2022). After dimensionality reduction, we visualised the data in a biplot along with environment vectors representing the microhabitat features onto the ordination. We used the *metaMDS* function of the ‘vegan’ package on the pooled abundance data from transects across seasons to determine if the composition of amphibians differed across land-use types. We used three-dimensional ordination with Bray-Curtis dissimilarity index for the non-metric multidimensional scaling (NMDS) analysis since the stress value for two-dimensional ordination was > 0.2. We tested for the homogeneity of multivariate dispersion using the *betadisper* function, and for the differences between the land-use types using the permutational multivariate analysis of variance (PERMANOVA) as implemented by the *adonis* function in package ‘vegan’.

We used a generalised linear mixed model with a negative-binomial error structure to determine the differences in the overall amphibian abundance across the different land-use types. We used a nested random effect structure in the instances in which the intercept varied among plateaus and transects within plateaus. We chose this structure after assessing multiple competing random effect structures (including the ones with spatial coordinates; Table S3). We considered land-use types whose 95% CI on the estimated coefficients did not overlap zero to influence abundance significantly. We estimated marginal means and assessed pairwise contrasts for land-use types in the model using Tukey’s method for *p*-value adjustment. We estimated the marginal and conditional *R^2^* for the model. We used the R packages ‘glmmTMB’, ‘DHARMa’, ‘MuMIn’, ‘ape’, and ‘emmeans’ for this analysis (Bartoń, 2023; Brooks et al., 2017; Hartig, 2022; Lenth, 2023; Paradis and Schliep, 2019).

We assessed the alpha diversity of amphibians using Hill numbers (species richness, Hill-Shannon diversity, and Hill-Simpson diversity) on the individual-based abundance data using pooled data from all transects across temporal replicates. We generated interpolation and extrapolation sampling curves with bootstrapped confidence intervals using the package ‘iNEXT’ (Chao et al., 2014) for the three land-use types. To understand assemblage dissimilarity between localities across land-use types, we calculated *β*-diversity values for transects pooled across the temporal replicates. This analysis was carried out for both abundance and incidence data, as incidence- and abundance-based dissimilarity indices provide different information. While incidence-based beta diversity measures demonstrate species replacement and/or nestedness, abundance-based beta diversity measures may demonstrate whether abundances are similar across assemblages (balanced variation) or whether species abundances of some assemblages are subsets of others (abundance gradient) (Baselga, 2017). We used the ‘betapart’ package (Baselga and Orme, 2012) for computing abundance-based total multiple-site Bray-Curtis dissimilarities (*β_BC_*) and their components caused by balanced variation in abundance (*β_BC.BAL_*) and abundance gradients (*β_BC.GRA_*); and to compute incidence-based total multiple site Sørensen dissimilarities (*β_SOR_*), as well as their respective turnover (*β_SIM_*) and nestedness components (*β_SNE_*) (Baselga, 2017, 2010). To make dissimilarities comparable for strata with different numbers of transects, we computed *β* values by selecting nine transects from each land-use type and re-sampling ten times before calculating the abundance and incidence-based multiple-site dissimilarities (Baselga, 2010).

To understand how individual species respond to environmental covariates and how traits influence species responses, we used the hierarchical modelling of species communities framework (HMSC) (Ovaskainen et al., 2017). HMSC is a multivariate hierarchical generalised linear mixed modelling framework with Bayesian inference, which can accommodate random effects such as spatial arrangement and study design with community matrices. The analysis was conducted at the finest resolution of data (sub-plot) to control for potential variability across sub-plots in microhabitats. This dataset comprised nine species (see section 2.3), all 950 subplot temporal replications, and 1,274 individuals. We checked for multicollinearity before specifying environmental covariates. The final model included the following covariates: land-use class (plateau, orchard, paddy), rock-and shallow-pool volumes (cm^3^), maximum paddy depth (cm), stream cross-sectional volume (cm^3^), woody- and flush-vegetation cover (%), grass cover (%), surface water presence, elevation (m), monsoon phase (early: June–July, late: August– September), and the presence/absence of rain (during the beginning of sampling). We used the identity of subplots as random levels in the model and spatial arrangement of subplots (x-y coordinates) in the study design. To model the presence-absence of the amphibians, we used probit regression. We fitted the HMSC model with the ‘Hmsc’ package (Tikhonov et al., 2020), which included the 11 environmental covariates, four traits, and spatial arrangement of subplots as a random effect. We assumed the default prior distributions (Ovaskainen and Abrego, 2020) and sampled the posterior distribution with three Markov chain Monte Carlo (MCMC) chains. Each chain was run for 300,000 iterations, of which we removed the first 50,000 as burn-in, and the remaining ones were thinned by 1000 to yield 250 posterior samples per chain, resulting in 750 posterior samples in total. We assessed the MCMC convergence of the model using Gelman diagnostics and evaluated the explanatory power of the model using Tjur’s *R^2^*.

Using the variance partitioning approach, we assessed the relative contributions of different fixed and random effects in explaining the variation in species occurrence. For this analysis, we grouped the covariates into six broad categories - elevation, land use, monsoon phase, vegetation, water sources, and spatial random effect. We evaluated the influence of predictors on species occurrence by examining the ≥ 95% posterior probability on β parameters. Similarly, we evaluated the γ parameters, which correspond to the trait-environment relationship.

## 3. RESULTS

### 3.1 Microhabitat variations across land-use types

The PCoA demonstrated differences in microhabitat variation among land-use classes (Fig. 1a). The two main axes of PCoA collectively explained 36.7% of the variation in data. While the PCoA axis 1 was highly positively associated with grass cover and paddy depth, axis 2 was associated positively with flush vegetation and shallow pool volumes (Fig. 1b). Paddy was distinct from plateaus and orchards in terms of high grass cover and water depth. Orchards were distinct from plateaus and were characterised by woody vegetation and less rock pool volume. Orchards were distinct from paddy, with comparatively high rock pool volume. Plateaus were distinct from orchards and paddy in terms of high rock and shallow pool volumes, and flush vegetation.

**Figure 1.**
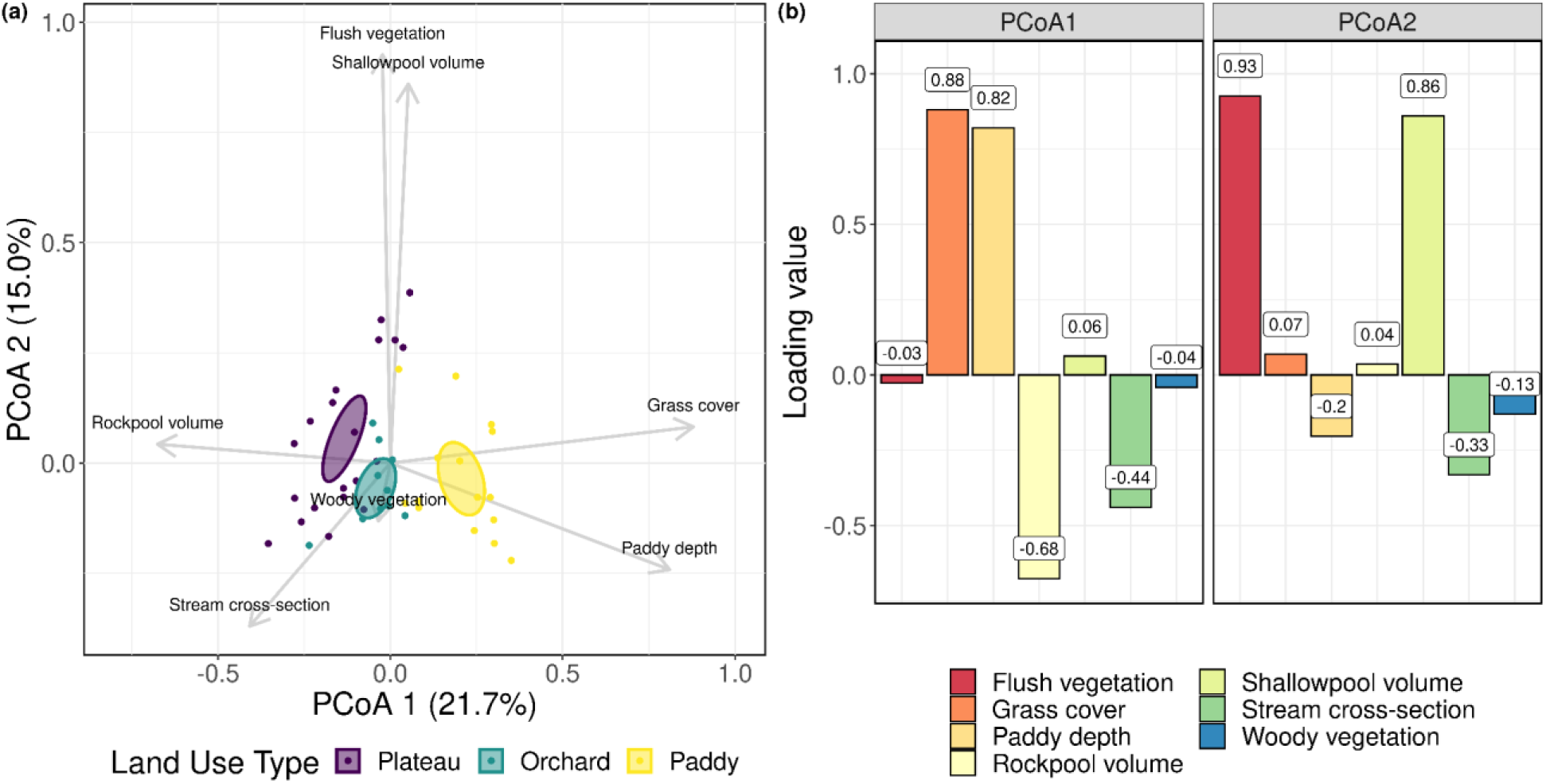
Principal coordinate analysis (PCoA) of the amphibian belt transects; (a) biplot showing the relationship among the transects (coloured dots) and habitat variables (grey vectors). Values in parentheses on x- and y-axes indicate the percentage of variation explained by each axis. Length of the vectors is proportional to the correlation between the variable and the PCoA ordination, and ellipses indicate multivariate 95% confidence intervals around the group centroid; (b) barplot showing the correlation of each habitat variable to the PCoA axes.

### 3.2 Amphibian community composition and abundance across land-use types

We encountered 1,279 individuals of 12 species of amphibians (Table S4) during the survey from four lateritic plateaus. These include nine genera from five families of Order Anura and one individual of *Gegeneophis* (Order: Gymnophiona). Among the 12 species, one is listed as ‘Endangered’, one is ‘Data Deficient’ in the IUCN RedList, and six are endemic to the Western Ghats. The most numerically dominant family was Dicroglossidae (1151), followed by Microhylidae (70), Rhacophoridae (22), Ranidae (11), Bufonidae (3), and Grandisoniidae (1); of which nine species occurred in all land-use types, one species (*Uperodon mormoratus*) was only detected in paddy, two species (*Duttaphrynus melanostictus* and *Gegeneophis seshachari*) were only detected in plateaus, and orchards had no unique species. Amphibians were detected in all transects across the land-use types when temporal replicates were pooled. The number of detections ranged between 4 and 103 individuals per transect. The NMDS analysis showed that the amphibian communities composition differed between the three land-use types in at least two of the three dimensions (PERMANOVA *R^2^*= 0.27, df = 2, *F* = 7.74, *p* = 0.001; Fig. 2a).

**Figure 2.**
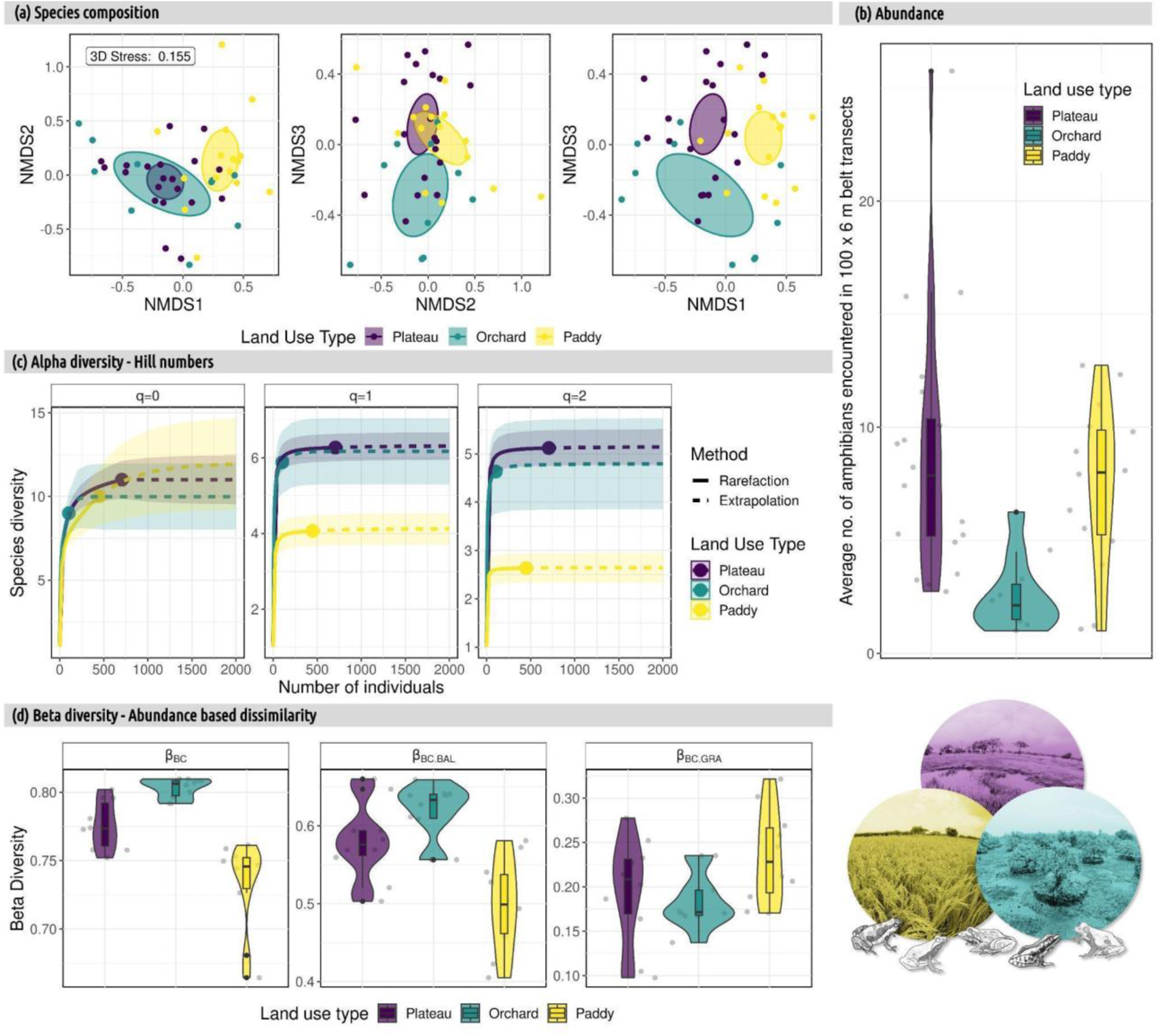
(a) Non-metric multidimensional scaling in three dimensions showing dissimilarities in the taxonomic composition between the three land-use types of amphibians in the lateritic plateaus of the northern Western Ghats. The ellipses indicate multivariate 95% confidence intervals around the group centroid; (b) violin plots showing the distribution of the number of individuals of amphibians seen across different transects; and (c) rarefaction-extrapolation curves showing amphibian species diversity by the number of individuals sampled across the land-use types for Hill numbers representing species richness (q = 0), Hill-Shannon (q = 1), and Hill-Simpson (q = 2) diversity indices. The shaded area corresponds to the 95% confidence interval; (d) violin plots showing the distribution dissimilarity indices across the three land-use types for abundance-based dissimilarity, where β_BC_ is total Bray-Curtis dissimilarity, β_BC.BAL_ is the component of dissimilarity due to balanced variation in abundance (analogous to turnover), and β_BC.GRA_ is the component of dissimilarity due to abundance gradients (analogous to nestedness). The grey dots are individual data points. Photographs and Illustrations by V. Jithin.

Land-use type explained significant variation in abundances (Nakagawa’s *R^2^* = 0.26). The contrast analysis showed that the abundance in orchards was significantly lower than that in plateaus and paddy. However, paddy did not differ significantly from plateaus (Fig. 2b; Table S5). The rank-abundance curves showed fewer frogs in orchards. The three *Minervarya* species had intermediate abundance in plateaus, whereas only *Minervarya syhadrensis* dominated the paddy and the rest were rare (Fig. S2).

### 3.3 Amphibian diversity across land-use types

The overall species richness (q = 0; *α*-diversity) patterns (which includes the rare species) did not significantly differ between land-use types as inferred from overlapping 95% CI. Hill-Shannon (q = 1) (representing common species) and Hill-Simpson diversity (q = 2) (representing dominant species) were lower in paddy than orchards and plateau, both of which had similar values of Hill-Shannon and Hill-Simpson (Fig. 2c).

Abundance-based Bray-Curtis indices (*β*-diversity) showed that the mean value of total dissimilarity (*β_BC_*) was highest in orchards (0.8), followed by the values in plateaus (0.78) and paddy (0.73) (Fig. 2d). The balanced variation component (*β_BC.BAL_*) also followed a similar trend (orchard = 0.62, plateau = 0.58, and paddy = 0.5), whereas the abundance gradient component (*β_BC.GRA_*) was higher in paddy (0.23), followed by that in plateaus (0.2) and orchards (0.18) (Fig. 2d). The incidence-based multiple site total Sørensen dissimilarity (*β_SOR_*) was higher in orchards (0.65), followed by the values in paddy (0.57) and plateaus (0.53). The turnover component (*β_SIM_*) was highest in orchards (0.49), followed by that in plateaus (0.39) and paddy (0.36), and nestedness component (*β_SNE_*) was highest in paddy (0.21), followed by that in orchards (0.16) and plateaus (0.14) (Fig. S3). Overall, paddy had the least *β-*diversity values, with a greater contribution of nestedness (*β_BC.GRA_*, *β_SNE_*), and orchards had the highest *β*-diversity values, with a greater contribution of replacement (*β_BC.BAL_*, *β_SIM_*).

### 3.4 Amphibian species-level response to land-use change

The average explanatory power (Tjur’s *R^2^*) of the HMSC model was 0.09. The *β* parameters showed that *Euphlycits jaladhara*, *Hoplobatrachus tigerinus, Minervarya cepfi*, and *Polypedates maculatus* showed a statistically supported negative response to orchards (Fig. 3a). While *M. syhadrensis* showed a positive response, *Minervarya gomantaki*, *Minervarya cepfi*, and *Euphlycits jaladhara* showed a negative response to paddy (Fig. 3a). Many species occurrences were positively associated with water-related covariates, such as rock-pool volume, paddy depth, and surface water presence (Fig. 3a). The explained variance attributed on average (across the species) was mainly explained by water sources (34.6%) followed by land-use type (18.4%) (Fig. 3b). No statistically supported trait-environmental relationships were found from the *γ* parameters (Fig. S4).

**Figure 3.**
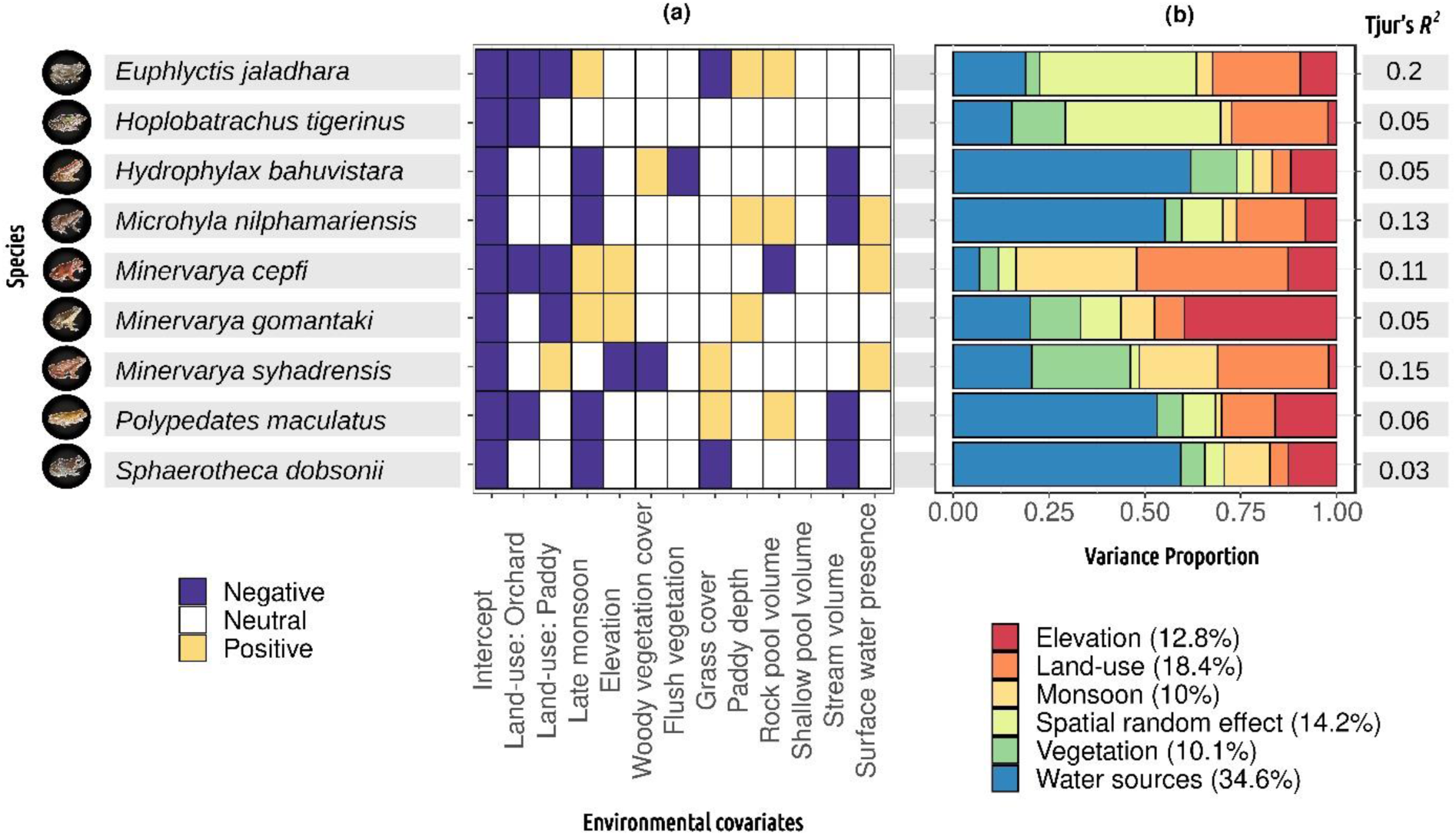
Hierarchical modelling of species communities model results showing (a) the mean posterior regression *β* parameter values measuring the species-specific responses of amphibians to each of the environmental covariates. Intercept represents the reference plateaus and early-monsoon. Violet colours indicate negative responses and mustard colours positive responses with ≥ 0.95 posterior probability; (b) the variance partitioning of explained variation among environmental covariates and random effect, with values in parentheses of legend markers showing the mean values across species. Illustrations by V. Jithin.

## 4. DISCUSSION

We examined how land use conversions associated with traditional paddy cultivation and recent agroforestry plantations impact rock outcrop frogs. Conversion to paddy and orchards differentially alters the availability of microhabitats for amphibians. Our assessments at the community- and species-level showed varied impacts of different forms of agriculture (paddy and orchards) on frogs, which is an important finding since most studies assess the combined impacts of different forms of agriculture. Following our expectation, conversion to paddy negatively affected the richness and homogenised the community as determined by more nested assemblages (community-level impacts), whereas conversion to orchards negatively impacted species persistence (species-level impacts). These results highlight the importance of examining community- and species-level impacts simultaneously to determine the varying impacts of land-use change on biodiversity. While past studies have assessed the impacts of land-use on forest-dwelling amphibians, to our knowledge, this is the first study that determines the impacts of land-use change on amphibians in threatened, open ecosystems.

Water resources, especially rock pools, paddy water depth and surface water, positively influenced multiple frog species. Large rock pools ensure that frog eggs and tadpoles are protected from desiccation during dry spells in the middle of the monsoon (see Fig. S5), making them critical microhabitats for maintaining amphibian populations on the plateau. Importance of seasonal and semi-permanent wetlands in maintaining and restoring amphibian populations has been shown in forested landscape previously (Karraker & Gibbs, 2009). In open ecosystems, such as deserts and low-elevation lateritic plateaus, rock pools are known as critical habitats for biodiversity, including amphibians (Thorpe et al., 2018; Vale et al., 2015). In such open habitats exposed to extreme climate, microhabitats such as pools and loose rocks (Jithin et al., 2023) can serve as an oasis harbouring biodiversity. In light of climate change, when the temperatures are expected to rise and the rainfall patterns change in the study area (Todmal, 2021), large rock pools will play a critical role for ensuring persistence of amphibians in these open ecosystems. Future studies on larval ecology of amphibians on the outcrops can throw more light on the value of rock pools for frogs. The plateaus are getting converted at an alarming rate to orchards (Bhattacharyya et al., 2019), and this study demonstrates the need to maintain or create large rock pools to enable amphibians to persist in the landscape.

Land-use was an important driver influencing amphibian prevalence. While most studies focus on determining the impacts of land-use change on diversity or species responses, the proximate drivers of change (e.g., loss of microhabitats) are infrequently documented (but see Barrios & Mello, 2022; Sueyoshi et al., 2016; Wood et al., 2017). Microhabitat changes can impact adult frog dispersion, larval occurrence, and morphology (Marques et al., 2018). Our study demonstrates that the conversion of plateaus to paddy and orchards significantly alters the microhabitat availability for amphibians. While paddy homogenised the habitat due to submergence, orchards reduce the availability of microhabitats, particularly rock pools, which are critical microhabitats for amphibians. Given the rapid expansion of orchards on lateritic plateaus, engagement with land owners is required to determine strategies to retain these vital microhabitats in orchards. Future studies need to also evaluate the efficacy of such measures in retaining frog populations.

Studies often pool data from different forms of agriculture when comparing impacts of agriculture on biodiversity (Gonthier et al., 2014; Newbold et al., 2015). At the species- and community-level, our study shows varying impacts of different forms of agriculture (paddy and orchards) on frogs. At the species-level, orchards negatively impacted four of the nine frog species. Moreover, the overall abundance of frogs in orchards was significantly lower than that in paddy and plateaus. Thus, microhabitat loss may be responsible for the reduced prevalence and abundance of frogs in orchards. All the frog species in our study site prefer stagnant pools for breeding. Paddy offers deeper stagnant pools to frogs throughout the season than plateaus. The deeper pools of paddy can protect the eggs/tadpoles from desiccation during the dry spells. Despite this, three species (two endemic and one range-restricted species) of frogs were negatively impacted by paddy. There is a need for a better understanding of the natural history of these frogs that can help us understand the differential responses of species to land-use change.

At the community-level, we found that paddy had lower richness of common and dominant species and had more nested communities than plateaus and orchards. This is indicative of biotic homogenisation as a likely consequence of habitat homogenisation. Interestingly, the orchards had similar richness as plateaus but higher turnover than plateaus; this is a likely consequence of rarity of frogs in orchards as has also been reported elsewhere (Jamoneau et al., 2017). If we had examined patterns only at species- or community-level instead of both, our conclusions could have been different. Examining at both species- and community-level gave us a holistic perspective of land-use change impact on amphibians. Given the variable impacts of different forms of agriculture on biodiversity, it is important to identify the probable mechanisms through which different forms of agriculture impact biodiversity. This will provide insights into suitable modifications that are required in existing land management practices to make these modified landscapes biodiversity-friendly. In this study, instead of comparing only frog diversity or species responses across land-use types, we also compared changes in microhabitat availability and species-microhabitat relationships, which enabled us to determine the important role of microhabitats, such as rock pools for frogs, and the reduced availability of such microhabitats in modified habitats.

*M. syhadrensis*, a species that is widely found in South Asia occurred more commonly in paddy than in the reference plateau ecosystems. Land-use change is known to benefit certain species, while negatively impacting others (McKinney & Lockwood, 1999). Conversion to paddy positively impacted one species of *Minervarya* (*M. syhadrensis*) while negatively impacting two other closely-related northern Western Ghats endemic species (*M. cepfi* and *M*. *gomantaki*). Replacement of specialists by generalists may lower the effect sizes of diversity differences across land-use types, but it results in loss of specialists that are often threatened or endemic species. This is of conservation concern, and further highlights the need to evaluate species-level responses, particularly in regions that harbour species of conservation importance.

None of the low-elevation lateritic plateaus are included within the protected area network and, unfortunately, are classified as ‘wastelands’. Historically, people have depended on these outcrops as pastures for their cattle and for growing paddy. Economic and other constraints have led to a reduction in paddy cultivation on the plateau. However, there has been a drastic expansion of orchards on the plateaus. Unlike paddy, mango and cashew are cash crops and in good years, the local alphonso mangoes grown on plateaus fetch good profits for farmers. Farmers from the region have been awarded ‘Global Good Agricultural Practises’ which facilitates mango exports globally. More than 150,000 ha of land in the region has been converted to mango orchards, a significant proportion of which is on the lateritic plateaus. Areas with extensive lateritic plateaus have been identified as ‘highly suitable’ areas for mango cultivation (Salunkhe et al., 2023). Moreover, many orchards are fenced off with rocks (Fig. S6), which may prevent movement of terrestrial amphibians on the plateau, an aspect that needs closer examination. Additionally, the plateaus have been identified for other development activities, including oil refinery (Deshpande, 2023). There is a need for systematic surveys to identify priority plateaus for conservation. Since most of these rock outcrop areas are not legally protected and are within privately owned land, community-based conservation initiatives, focussed on the ecological and sociological needs of the community are required. It is critical to engage with the private landowners to ensure that while their private lands are converted to orchards, critical microhabitats, such as pools, are retained. There is an urgent need to make necessary amendments in the government policy so that these unique ecosystems are not classified as wastelands.

### 4.1. Conservation Implications

Results of our study have broader implications for global assessments of biodiversity responses to anthropogenic changes, in which often different types of agricultural land-use types or management intensity are merged together to arrive at general conclusions (Tuck et al., 2014). Our study demonstrates the need to evaluate the impacts of different forms of agriculture separately as different forms of agriculture can vary in their intensity and nature of impact. The study also has a recommendation for researchers and managers relying on single measures of biodiversity (Duelli and Obrist, 2003) that it is critical to determine community- and species-level impacts in parallel as they capture different information which may often be delinked from each other. In seemingly barren ecosystems, such as rock outcrops, which are going to be exposed to extreme climatic events, it is critical to identify key microhabitats that help sustain biodiversity. There is a need to recognize the unique ecological value of the unprotected rock outcrops, prioritise sites for conservation, and protect them from land-use change in partnership with local communities.

## ACKNOWLEDGEMENTS

We thank the Maharashtra Forest Department, particularly the Chief Wildlife Warden, Sunil Limaye, for giving us the necessary permits (Letter No. Desk-22(8)/WL/Research/CR-53(20-21)/3361/22-22) to conduct the study. We thank On the Edge (UK), The Bombay Environmental Action Group, and The Habitat Trust (India) for funding this work. We thank Varad B. Giri for useful discussions, Mihir Kulkarni for running the HMSC analysis at the Centre for Cellular and Molecular Biology, and Saunak P. Pal for measuring the traits of specimens at the Bombay Natural History Society. We thank Aditya Gadkari, Ajay Nachinekar, Chandrakanth Gurav, Harshad Tulpule, Kamalakar Gurav, Pooja Ghate, Poorva Joshi, Pradeep Dingankar, Rakesh Patil, Rashmi Karandikar, Ravindra Karandikar, Santosh Padhye, Shailesh Joshi, Suhas Gurjar, Sujan Dandekar, and Yash Vichare for support during fieldwork. RN thanks Anand Osuri, Kulbhushansingh Suryawanshi, Suri Venkatachalam, and Madhura Niphadkar; and VJ thanks Abhijit Das, Jahnavi Joshi, Vijay Karthick, Nayantara Biswas, and Yukti Taneja for useful discussions. We thank Karthik T. for proofreading the manuscript.

## AUTHORS’ CONTRIBUTIONS

RN and VJ conceived the ideas and designed the methodology with inputs from AW; VJ and MR collected the data; VJ and RN analysed the data; VJ and RN led the writing of the manuscript with inputs from AW and MR. All authors contributed critically to the drafts and gave final approval for publication.

## DATA AVAILABILITY STATEMENT

Data and codes used in this study will be uploaded on DataDryad on acceptance.

## Notes

### Competing Interest Statement

The authors have declared no competing interest.

### Summary of Updates

Updated the Figure 3a (HMSC Beta plot) to correct an error in the analysis. Main text (Results and Discussions) updated accordingly. Updated the acknowledgement section.

## REFERENCES

Asad, S., Abrams, J.F., Guharajan, R., Lagan, P., Kissing, J., et al., 2021. Amphibian responses to conventional and reduced impact logging. For. Ecol. Manag. 484, 118949.

Barrios, M., Mello, F.T. de, 2022. Urbanization impacts water quality and the use of microhabitats by fish in subtropical agricultural streams. Environ. Conserv. 49, 155–163.

Bartoń, K., 2023. Multi-model inference.

Baselga, A., 2017. Partitioning abundance-based multiple-site dissimilarity into components: Balanced variation in abundance and abundance gradients. Methods Ecol. Evol. 8, 799– 808.

Baselga, A., 2013. Separating the two components of abundance-based dissimilarity: balanced changes in abundance vs. abundance gradients. Methods Ecol. Evol. 4, 552–557.

Baselga, A., 2010. Partitioning the turnover and nestedness components of beta diversity. Glob. Ecol. Biogeogr. 19, 134–143.

Baselga, A., Orme, C.D.L., 2012. betapart: an R package for the study of beta diversity. Methods Ecol. Evol. 3, 808–812.

Bhagwat, S.A., Willis, K.J., Birks, H.J.B., Whittaker, R.J., 2008. Agroforestry: a refuge for tropical biodiversity? Trends Ecol. Evol. 23, 261–267.

Bhattacharyya, T., Salvi, B.R., Haldankar, P.M., Salvi, N.V., 2019. Growing Alphonso Mango on Konkan Laterites, Maharashtra. Indian J. Fertil. 15, 878–885.

Bond, W.J., 2019. Open ecosystems: ecology and evolution beyond the forest edge. Oxford University Press.

Brooks, M.E., Kristensen, K., Benthem, K.J. van, Magnusson, et al., 2017. glmmTMB Balances Speed and Flexibility Among Packages for Zero-inflated Generalized Linear Mixed Modeling. R J. 9, 378–400.

Chao, A., Gotelli, N.J., Hsieh, T.C., Sander, E.L., Ma, K.H., Colwell, R.K., Ellison, A.M., 2014. Rarefaction and extrapolation with Hill numbers: a framework for sampling and estimation in species diversity studies. Ecol. Monogr. 84, 45–67.

Cingolani, A.M., Cabido, M., Gurvich, D.E., Renison, D., Díaz, S., 2007. Filtering processes in the assembly of plant communities: Are species presence and abundance driven by the same traits? J. Veg. Sci. 18, 911–920.

Cordier, J.M., Aguilar, R., Lescano, J.N., Leynaud, G.C., Bonino, A., et al., 2021. A global assessment of amphibian and reptile responses to land-use changes. Biol. Conserv. 253, 108863.

DeFries, R., Rosenzweig, C., 2010. Toward a whole-landscape approach for sustainable land use in the tropics. Proc. Natl. Acad. Sci. 107, 19627–19632.

Deshpande, A., 2023. An oil refinery in Maharashtra is dividing villages in the Konkan belt. The Hindu. (Accessed on 15 September 2023). https://www.thehindu.com/news/national/an-oil-refinery-in-maharashtra-is-dividing-villages-in-the-konkan-belt/article66821061.ece

Duelli, P., Obrist, M.K., 2003. Biodiversity indicators: the choice of values and measures. Agric. Ecosyst. Environ., 98, 87–98.

Fulgence, T.R., Martin, D.A., Randriamanantena, R., Botra, R., Befidimanana, E., et al., 2022. Differential responses of amphibians and reptiles to land-use change in the biodiversity hotspot of north-eastern Madagascar. Anim. Conserv. 25, 492–507.

Ganeshmurthy, A.N., Rupa, T.R., Shivananda, T.N., 2018. Enhancing Mango Productivity through Sustainable Resource Management. J. Hortic. Sci. 13, 1–31.

Garg, S., Biju, S.D., 2017. Description of four new species of Burrowing Frogs in the Fejervarya rufescens complex (Dicroglossidae) with notes on morphological affinities of Fejervarya species in the Western Ghats. Zootaxa 4277, 451–490.

Gonthier, D.J., Ennis, K.K., Farinas, S., Hsieh, H.-Y., Iverson, A.L., et al., 2014. Biodiversity conservation in agriculture requires a multi-scale approach. Proc. R. Soc. B Biol. Sci. 281, 20141358.

Hartig, F., 2022. DHARMa: Residual Diagnostics for Hierarchical (Multi-Level / Mixed) Regression Models.

Hof, C., Araújo, M.B., Jetz, W., Rahbek, C., 2011. Additive threats from pathogens, climate and land-use change for global amphibian diversity. Nature 480, 516–519.

Jamoneau, A., Passy, S. I., Soininen, J., Leboucher, T., Tison-Rosebery, J., 2018. Beta diversity of diatom species and ecological guilds: Response to environmental and spatial mechanisms along the stream watercourse. Freshw. Biol., 63, 62–73.

Jesse, W.A.M., Behm, J.E., Helmus, M.R., Ellers, J., 2018. Human land use promotes the abundance and diversity of exotic species on Caribbean islands. Glob. Change Biol. 24,

Jithin, V., Rane, M., Watve, A., Giri, V.B., Naniwadekar, R., 2023. Between a rock and a hard place: Comparing rock-dwelling animal prevalence across abandoned paddy, orchards, and rock outcrops in a biodiversity hotspot. Glob. Ecol. Conserv. e02582.

Kéfi, S., Domínguez-García, V., Donohue, I., Fontaine, C., Thébault, E., Dakos, V., 2019. Advancing our understanding of ecological stability. Ecol. Lett. 22, 1349–1356.

Komanduri, K.P.K., Sreedharan, G., Vasudevan, K., 2023. Abundance and composition of forest-dwelling anurans in cashew plantations in a tropical semi-evergreen forest landscape. Biotropica 55, 594–604.

Kulkarni, A., Shigwan, B.K., Vijayan, S., Watve, A., Karthick, B., Datar, M.N., 2022. Indian rock outcrops: review of flowering plant diversity, adaptations, floristic composition and endemism. Trop. Ecol.

Laurance, W.F., Sayer, J., Cassman, K.G., 2014. Agricultural expansion and its impacts on tropical nature. Trends Ecol. Evol. 29, 107–116.

Lenth, R., 2023. emmeans: Estimated Marginal Means, aka Least-Squares Means.

Madhusudan, M.D., Vanak, A.T., 2022. Mapping the distribution and extent of India’s semi-arid open natural ecosystems. J. Biogeogr. jbi.14471.

McKinney, M.L., Lockwood, J.L., 1999. Biotic homogenization: a few winners replacing many losers in the next mass extinction. Trends Ecol. Evol. 14, 450–453.

Myers, N., Mittermeier, R.A., Mittermeier, C.G., Da Fonseca, G.A., Kent, J., 2000. Biodiversity hotspots for conservation priorities. Nature 403, 853–858.

Newbold, T., Hudson, L.N., Hill, S.L., Contu, S., Lysenko, I., et al., 2015. Global effects of land use on local terrestrial biodiversity. Nature 520, 45–50.

Newbold, T., Oppenheimer, P., Etard, A., Williams, J.J., 2020. Tropical and Mediterranean biodiversity is disproportionately sensitive to land-use and climate change. Nat. Ecol. Evol. 4, 1630–1638.

Nori, J., Lemes, P., Urbina-Cardona, N., Baldo, D., Lescano, J., Loyola, R., 2015. Amphibian conservation, land-use changes and protected areas: A global overview. Biol. Conserv. 191, 367–374.

Nowakowski, A.J., Thompson, M.E., Donnelly, M.A., Todd, B.D., 2017. Amphibian sensitivity to habitat modification is associated with population trends and species traits: Nowakowski et al. Glob. Ecol. Biogeogr. 26, 700–712.

Oksanen, J., Simpson, G.L., Blanchet, F.G., Kindt, R., et al., 2022. vegan: Community Ecology Package.

Ovaskainen, O., Abrego, N., 2020. Joint species distribution modelling: with applications in R. Cambridge University Press.

Ovaskainen, O., Tikhonov, G., Norberg, A., Guillaume Blanchet, F., Duan, L., et al., 2017. How to make more out of community data? A conceptual framework and its implementation as models and software. Ecol. Lett. 20, 561–576.

Paradis, E., Schliep, K., 2019. ape 5.0: an environment for modern phylogenetics and evolutionary analyses in R. Bioinformatics 35, 526–528.

Parr, C.L., Lehmann, C.E.R., Bond, W.J., Hoffmann, W.A., Andersen, A.N., 2014. Tropical grassy biomes: misunderstood, neglected, and under threat. Trends Ecol. Evol. 29, 205– 213.

R Core Team, 2023. R: A Language and Environment for Statistical Computing.

Salunkhe, S., Nandgude, S., Tiwari, M., Bhange, H., Chavan, S.B., 2023. Land Suitability Planning for Sustainable Mango Production in Vulnerable Region Using Geospatial Multi-Criteria Decision Model. Sustainability 15, 2619.

Sankararaman, V., Dalvi, S., Miller, D.A., Karanth, K.K., 2021. Local and landscape characteristics shape amphibian communities across production landscapes in the Western Ghats. Ecol. Solut. Evid. 2, e12110.

Scott, N.J., Crump, M.L., Zimmerman, B.L., Jaeger, R.G., et al., 1994. Standard techniques for inventory and monitoring. Meas. Monit. Biol. Divers. Stand. Methods Amphib. W Ronald.

Smith, R.K., Meredith, H., Sutherland, W.J., 2020. Amphibian Conservation, in: What Works in Conservation 2020. Open Book Publishers, Cambridge, UK., pp. 9–64.

Solar, R.R. de C., Barlow, J., Ferreira, J., Berenguer, E., et al., 2015. How pervasive is biotic homogenization in human-modified tropical forest landscapes? Ecol. Lett. 18, 1108– 1118.

Sueyoshi, M., Ishiyama, N., Nakamura, F., 2016. β-diversity decline of aquatic insects at the microhabitat scale associated with agricultural land use. Landsc. Ecol. Eng. 12, 187–196.

Thorpe, C.J., Watve, A., 2015. Lateritic plateaus in the northern Western Ghats, India; a review of bauxite mining restoration practices. J. Ecol. Soc. 28, 25–44.

Tikhonov, G., Opedal, Ø.H., Abrego, N., Lehikoinen, A., de Jonge, M.M., et al., 2020. Joint species distribution modelling with the r-package Hmsc. Methods Ecol. Evol. 11, 442– 447.

Todmal, R.S., 2021. Future Climate Change Scenario over Maharashtra, Western India: Implications of the Regional Climate Model (REMO-2009) for the Understanding of Agricultural Vulnerability. Pure Appl. Geophys. 178, 155–168.

Tuck, S.L., Winqvist, C., Mota, F., Ahnström, J., et al., 2014. Land-use intensity and the effects of organic farming on biodiversity: a hierarchical meta-analysis. J. Appl. Ecol. 51, 746– 755.

Veldman, J.W., Overbeck, G.E., Negreiros, D., Mahy, G., Le Stradic, S., et al., 2015. Where Tree Planting and Forest Expansion are Bad for Biodiversity and Ecosystem Services. BioScience 65, 1011–1018.

Vié, J.-C., Hilton-Taylor, C., Stuart, S.N., 2009. Wildlife in a changing world: an analysis of the 2008 IUCN Red List of Threatened Species. IUCN.

Watve, A., 2013. Status review of Rocky plateaus in the northern Western Ghats and Konkan region of Maharashtra, India with recommendations for conservation and management. J. Threat. Taxa 5, 3935–3962.

Wood, J.R., Holdaway, R.J., Orwin, K.H., Morse, C., Bonner, K.I., et al., 2017. No single driver of biodiversity: divergent responses of multiple taxa across land use types. Ecosphere 8, e01997.

