## Supporting Files for "Orchards and paddy differentially impact rock outcrop amphibians: Insights from community- and species-level responses" (from another version of this preprint: version 1, 2023-10-05)

*This file includes:*

**TABLES**

**Table S1.** Sampling effort across different lateritic plateaus and land use types, along with the temporal replication details.

**Table S2.** Functional traits measured in the adult anurans of nine selected species.

**Table S3.** Multimodel inference table for generalised linear mixed models with negative-binomial error structure with varying random effects.

**Table S4.** Complete list of amphibians detected in the 100x6 m belt transects across the land-use types.

**Table S5.** The effect of land-use change on the overall amphibian abundance from the results of generalised linear mixed models with negative-binomial error structure.

**FIGURES**

**Figure S1.** Map showing transect locations across different land-use types on lateritic plateaus

**Figure S2.** Rank-Abundance curves indicating the species richness and evenness across the three land-use types.

**Figure S3.** Violin plots showing the distribution dissimilarity indices across the three land-use types for incidence-based dissimilarity.

**Figure S4.** Heatmap showing how species traits influence environmental responses.

**Figure S5.** Photographs showing dried rockpools and dead tadpoles.

**Figure S6.** Photographs showing rock walls in the orchards.

**TABLES**

**Table S1.** Sampling effort across different lateritic plateaus and land use types, along with the temporal replication details.

| **Land Use** | **Plateau Name** | **No. of 100 x 6 m transects** | **No. of temporal replications** | **No. of replicated transects** |
| --- | --- | --- | --- | --- |
| Plateau | Bakale | 5 | 4 | 20 |
|  | Devache Gothane | 5 | 4 | 20 |
|  | Devi Hasol | 5 | 4 | 20 |
|  | Gaonkhadi | 5 | 4 | 20 |
| Orchard | Devache Gothane | 5 | 2 | 10 |
|  | Devi Hasol | 5 | 4 | 20 |
|  | Gaonkhadi | 5 | 4 | 20 |
| Paddy | Devache Gothane | 5 | 4 | 20 |
|  | Devi Hasol | 5 | 4 | 20 |
|  | Gaonkhadi | 5 | 4 | 20 |
| **Total** | | **50** | **38** | **190** |

**Table S2.** Functional traits measured in the adult anurans of nine selected species. Highly correlated traits, and those explained little variation among the species were removed (as mentioned in the remarks column). Values for the selected four traits (SVL, relative hindlimb length, relative eye size and eye position were scaled and centred before the analysis. Only measurements (in mm) of male specimens were used in the analysis to ensure consistency across species, with available information.

| **Functional Trait** | **Ecological Relevance** | **Description/ Data Type** | **Reference** | **Remarks** |
| --- | --- | --- | --- | --- |
| **Body Size (SVL)** | Related to heat and water balance; nutrient cycling and energy flow through trophic webs as prey and predator | Continuous variable | Hernández-Ordóñez et al., 2019; Olalla-Tárraga et al., 2009 |  |
| **Toe** **Webbing** | Related to habitat; extended webbing is found in aquatic species compared to arboreal and terrestrial ones. Also related to thermoregulation, hydroregulation and protection against predation. | Increasing degree of toe webbing was considered as: basal, small, medium, large (Garg & Biju, 2017). This was converted to continuous variable by scoring it as 1-4 | Cortes-Gomez et al., 2015; Dehling & Dehling, 2021; Hernández-Ordóñez et al., 2019 | Highly correlated (r=0.9) with the Relative HIndlimb Length. Removed from subsequent analysis. |
| **Head** **Shape** | Related to foraging strategy - habitat and microhabitat use, and feeding biology | Continuous variable. Calculated by Head Length/ Head Width | Dalmolin et al., 2020; Paluh et al., 2020 | Highly correlated (r=0.74) with the Body Size. Removed from subsequent analysis. |
| **Relative Hindlimb Length** | Related to dispersion ability, and indicative of habitat/ strata | Continuous variable. Calculated by [Femur+Shank Length]/ SVL | Dehling & Dehling, 2021; Hernández-Ordóñez et al., 2019; Pittman et al., 2008 |  |
| **Egg Deposition Type** | Related to breeding habitat - direct/ in water/ on surface without foam cover/ foam nest/ eggs deposited within a foam nest - Embryo insulation from temperature fluctuation | Categorical variable with levels: foam nesting/ in water: converted to continuous by marking 1 and 0 | Dehling & Dehling, 2021 | Not used in the analysis, as only one species (*Polypedates*) was using a foam nest. All others were using water. |
| **Eye Position** | Associated with habitat exploration capabilities, prey detection (foraging), partner evaluation (reproduction), and predator avoidance | Continuous variable. Calculated by Interorbital Distance/ Head Width | Dalmolin et al., 2020 |  |
| **Relative Eye Size** | Same as above | Continuous variable. Calculated by Eye Diameter/ SVL | Dalmolin et al., 2020 |  |

**Table S3.** Multimodel inference table for generalised linear mixed models with negative-binomial error structure with varying random effects. All models contained the same fixed effect (land-use type) and the response variable was overall amphibian abundance.

| **Model Name** | **Random effect structure** | **df** | **Log Likelihood** | **AICc** | **ΔAICc** | **Weight** |
| --- | --- | --- | --- | --- | --- | --- |
| **Model 2** | Random intercept varying among plateaus and among transects within plateaus  (nested random effect) | 6 | -542.18 | 1096.8 | 0 | 0.63 |
| **Model 3** | Random intercept varying among transects; spatial autocorrelation* accounted using spatial coordinate Matérn | 8 | -541.33 | 1099.5 | 2.63 | 0.17 |
| **Model 1** | Random intercept varying among transects | 5 | -544.87 | 1100.1 | 3.24 | 0.12 |
| **Model 4** | Nested random effect; spatial autocorrelation accounted | 9 | -540.96 | 1100.9 | 4.08 | 0.08 |

*Temporal replicates one and four (June and September) showed significant positive spatial autocorrelation (p<0.001).

**Table S4.** Complete list of amphibians detected in the 100 × 6 m^2^ belt transects across the land-use types with overall number of detections, endemism and IUCN status. IUCN status abbreviated as NA - Not Assessed, LC - Least Concern, EN - Endangered, DD - Data Deficient,

| **Family** | **Species** | **Endemism/ IUCN Status** | **Number of detections** | | | |
| --- | --- | --- | --- | --- | --- | --- |
|  |  |  | **Plateau** | **Orchard** | **Paddy** | **Total** |
| Bufonidae | Common Asian Toad (*Duttaphrynus melanostictus)* | LC | 3 | 0 | 0 | **3** |
| Dicroglossidae | Jaladhara Skittering Frog (*Euphlyctis jaladhara*) | Peninsular India (NA) | 231 | 12 | 39 | **282** |
|  | Indian Bull Frog (*Hoplobatrachus tigerinus*) | LC | 69 | 17 | 44 | **130** |
|  | CEPF Burrowing Frog (*Minervarya cepfi*) | Northern Western Ghats (NA) | 115 | 7 | 12 | **134** |
|  | Goan Fejervarya (*Minervarya gomantaki*) | Northern Western Ghats (NA) | 108 | 39 | 40 | **187** |
|  | Bombay Wart Frog (*Minervarya syhadrensis*) | Western Ghats (LC) | 113 | 24 | 264 | **401** |
|  | Dobson's Burrowing Frog (*Sphaerotheca dobsonii*) | Peninsular India (LC) | 15 | 12 | 2 | **29** |
| Grandisoniidae | Seshachari's Caecilian (*Gegeneophis seshachari*) | Northern Western Ghats (DD) | 1 | 0 | 0 | **1** |
| Microhylidae | Nilphamarai Narrow-mouthed Frog (*Microhyla nilphamariensis*) | Peninsular India (NA) | 34 | 3 | 34 | **71** |
|  | Mottled Globular Frog (*Uperodon mormoratus*) | Western Ghats (EN) | 0 | 0 | 1 | **1** |
| Ranidae | Wide-spread Fungoid Frog (*Hydrophylax bahuvistara*) | Western Ghats (NA) | 7 | 9 | 1 | **17** |
| Rhacophoridae | Common Indian Treefrog (*Polypedates maculatus*) | LC | 10 | 2 | 11 | **23** |
| **Total** | | | **706** | **125** | **448** | **1279** |

**Table S5** **(a).** The effect of land-use change on the overall amphibian abundance from the results of generalised linear mixed models with negative-binomial error structure. Rows highlighted in bold indicate coefficients whose 95% CI do not overlap zero and thus were inferred to significantly influence the response variable. *Marginal R^2^ / Conditional R^2^*= *0.264 / 0.458*.

| ***Random Effects***  *[N_transect_=50; N_plateau_=4; No. of observations=190; Dispersion parameter for negative binomial family=1.74; σ^2^=0.57; ICC=0.26]* | | | |
| --- | --- | --- | --- |
| **Groups** | **Variance (***τ_00_***)** | **Standard Deviation** | **95% CI** |
| Transect: Plateau | 0.08 | 0.28 | 0.13 – 0.63 |
| Plateau | 0.12 | 0.35 | 0.12 – 1.01 |
| ***Fixed Effects*** | | | |
| **Term** | **Estimate** | **95% CI** | ***p*** |
| **Intercept: Plateau** | **2.06** | **1.65 – 2.47** | **< 0.001** |
| **Orchard** | **-1.27** | **-1.68 – -0.87** | **< 0.001** |
| Paddy | -0.26 | -0.62 – 0.11 | 0.17 |

**S5 (b).** Estimated marginal means for land-use combinations in negative binomial GLMM assessing the effect of land-use change on the overall abundance of amphibians. Rows highlighted in bold indicate estimated marginal means whose 95% CI do not overlap zero and thus were inferred to significantly influence the response variable. Results are given on the log (not the response) scale. *p*-values were adjusted using Tukey’s method.

| **Contrast** | **Estimate** | **SE** | ***z*** | ***p*** |
| --- | --- | --- | --- | --- |
| **Plateau – Orchard** | **1.27** | **0.21** | **6.14** | **< 0.0001** |
| Plateau – Paddy | 0.26 | 0.19 | 1.37 | 0.36 |
| **Orchard – Paddy** | **-1.02** | **0.21** | **-4.8** | **< 0.0001** |

**FIGURES**

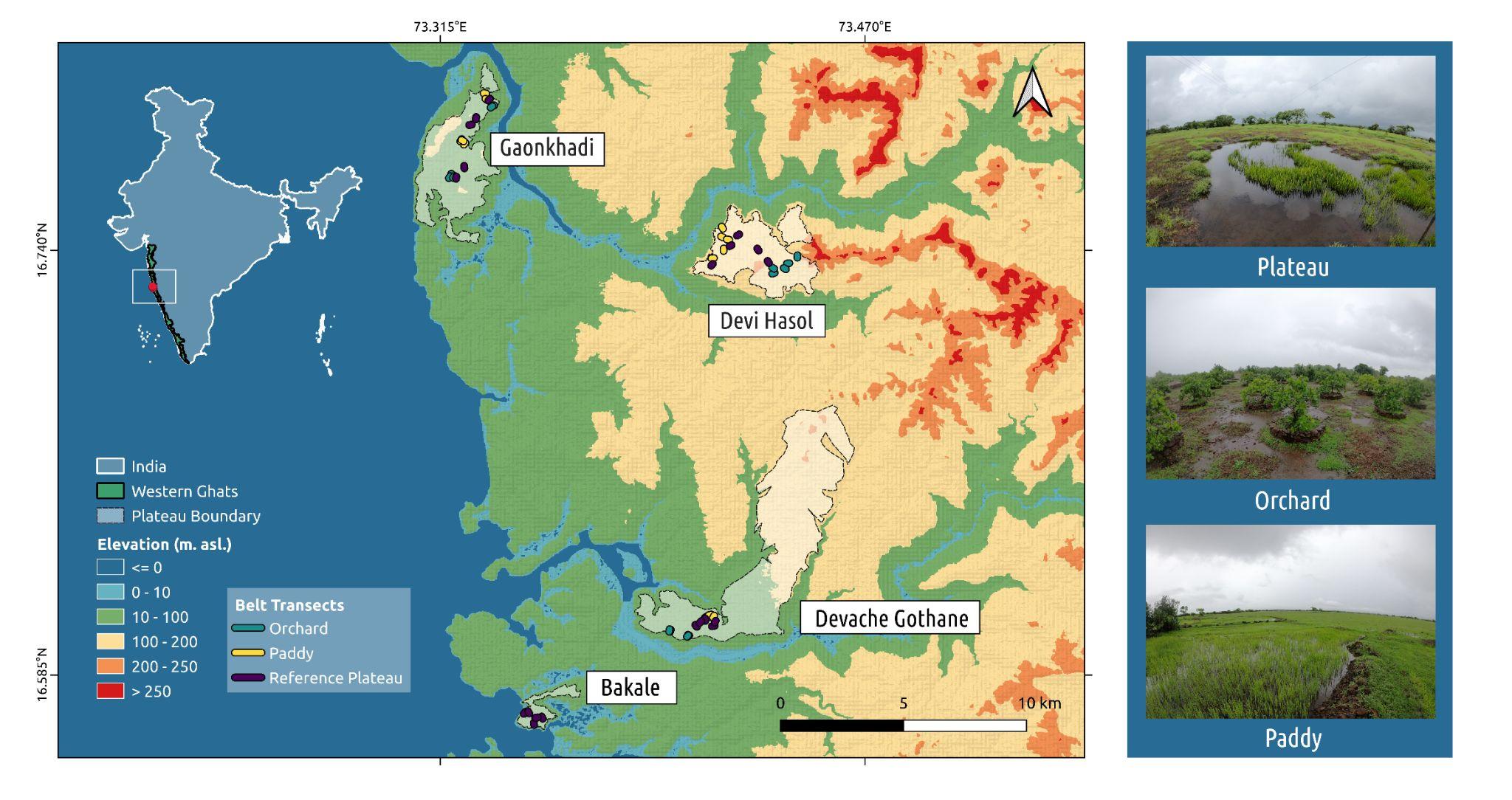

**Figure S1.** Map showing transect locations across different land-use types on lateritic plateaus (shown in the polygons) in the northern Western Ghats region of Ratnagiri, Maharashtra, India. Photographs by V. Jithin.

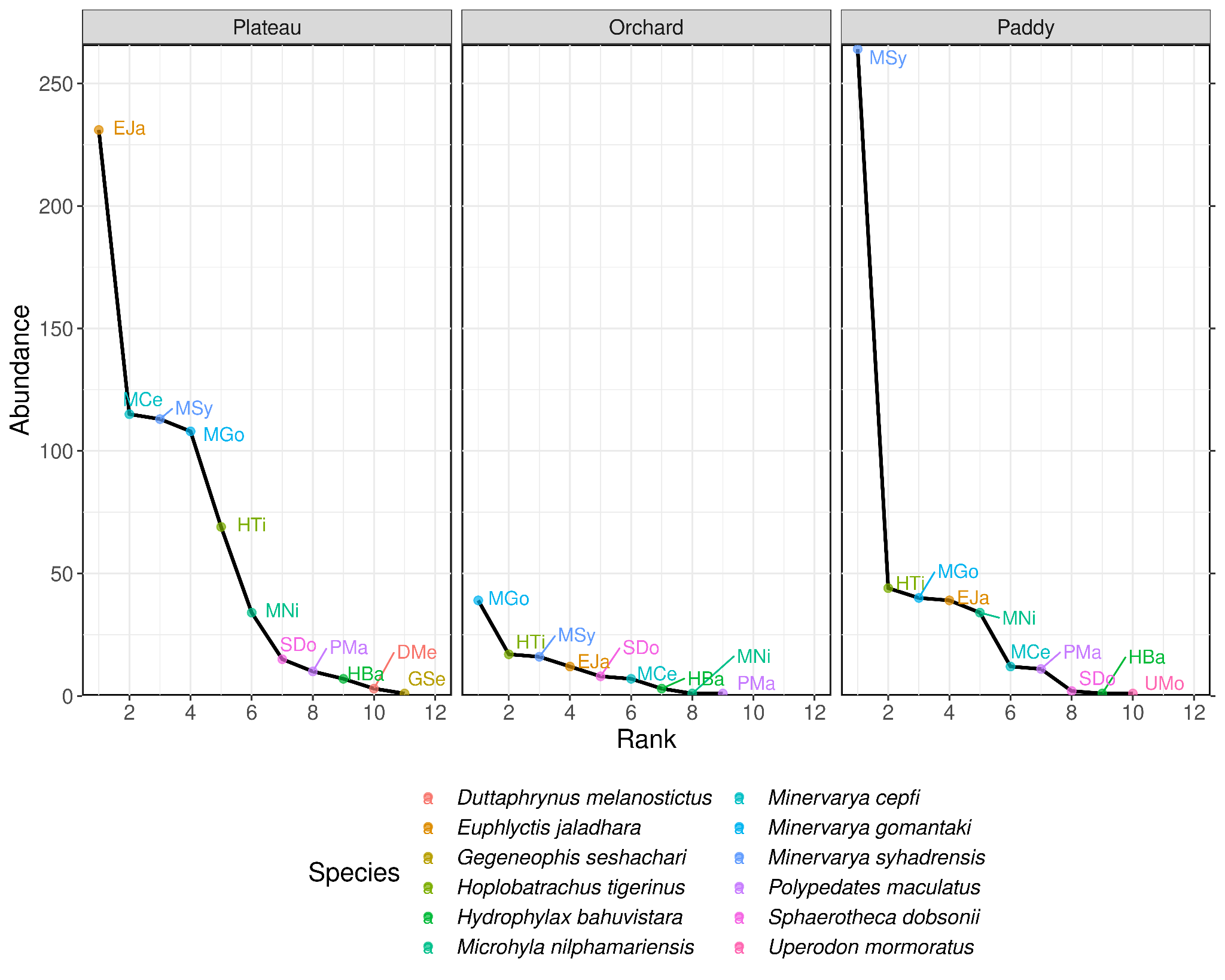

**Figure S2.** Rank-Abundance curves indicating the species richness and evenness across the three land-use types. Abbreviations in the plot are colour coded to the legend.

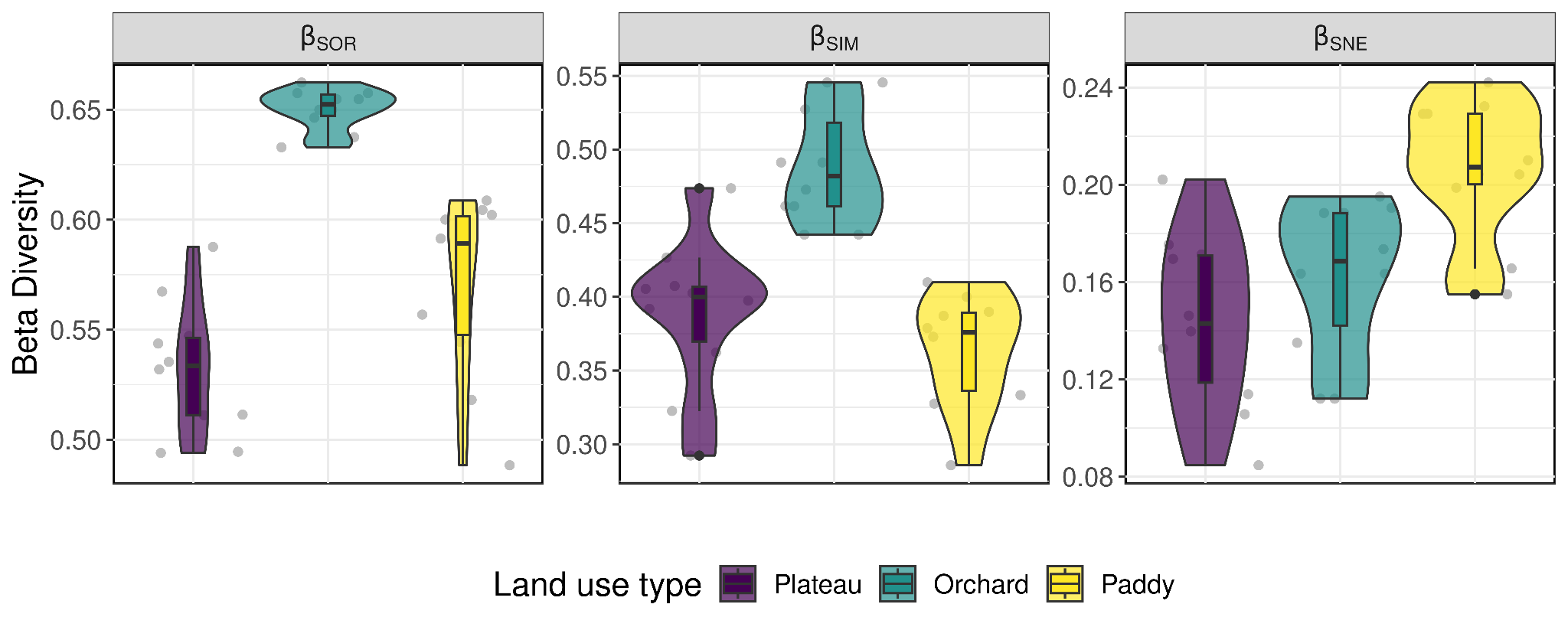

**Figure S3.** Violin plots showing the distribution dissimilarity indices across the three land-use types for incidence-based dissimilarity, where β_SOR_ is total Sørensen dissimilarity, β_SIM_ is the component of dissimilarity due to species replacement, and β_SNE_ is the component of due to nestedness. The grey dots are individual data points.

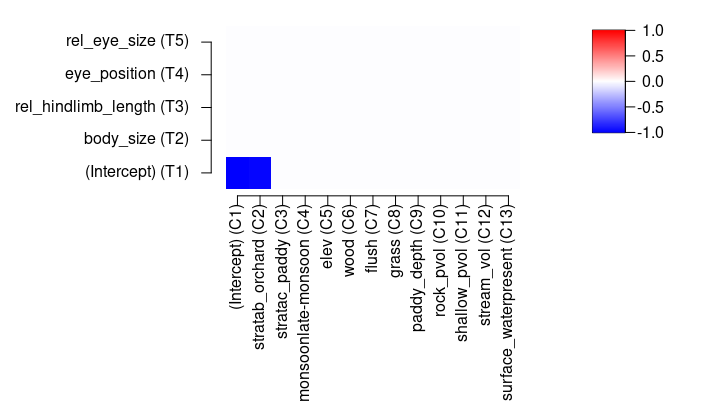

**Figure S4.** Heatmap showing how species traits influence environmental responses in the case of rock outcrop amphibians in the northern Western Ghats. Intercept: Paddy; stratab_orchard: Land use - Orchard; stratac_paddy: Land use - Paddy; monsoonlate-monsoon: Late-monsoon; elev: Elevation; wood: Woody vegetation (%); flush: Flush vegetation (%); grass: Grass Cover (%); paddy_depth: Paddy water depth; rock_pvol: Rock pool volume; shallow_pvol: Shallow pool volume; stream_vol: Stream cross-sectional volume, surface_waterpresent: Presence of surface water

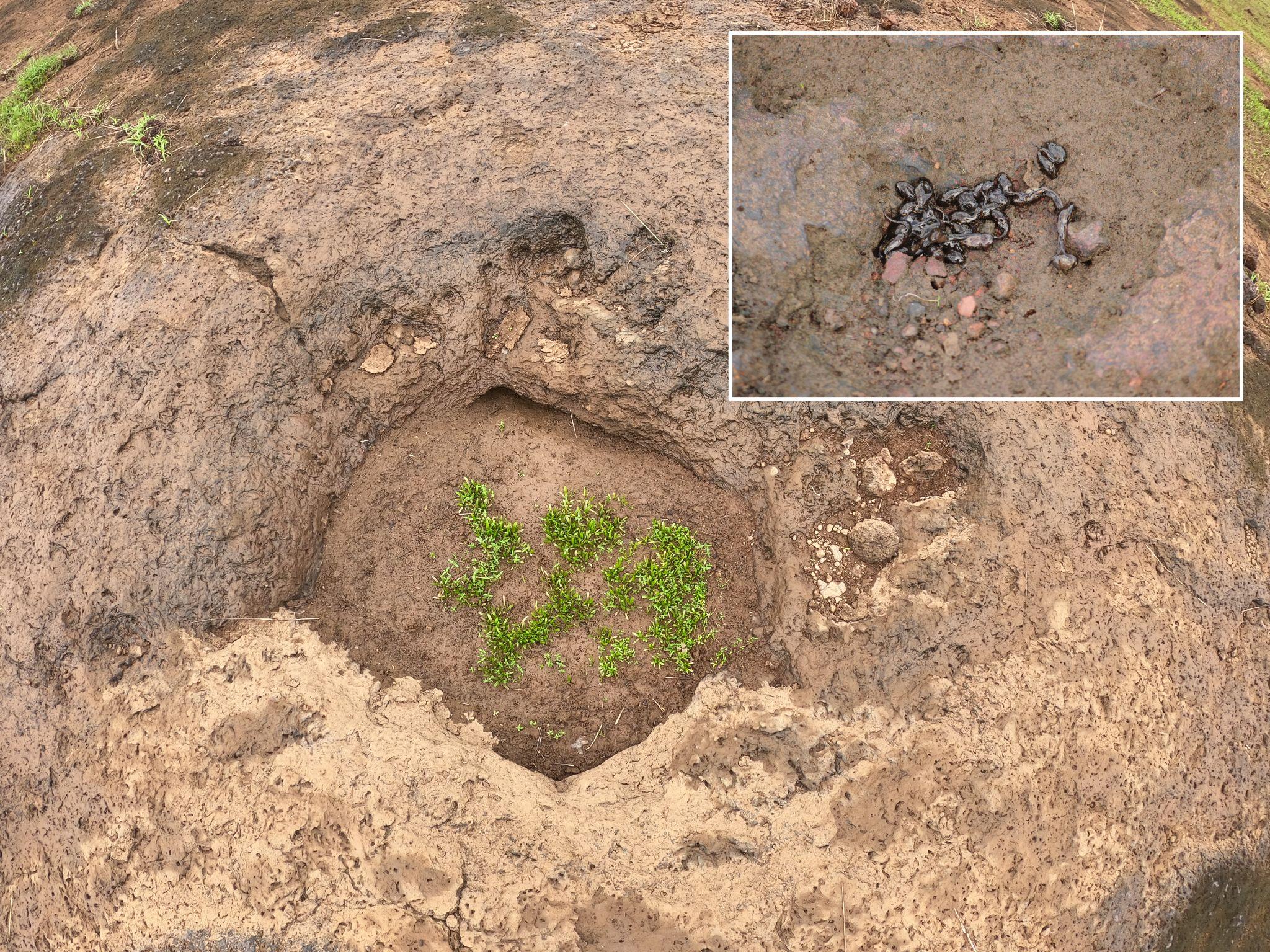

**Figure S5.** A rockpool dried due to irregular patterns in rain. The inset image shows the dead tadpoles. Photographs: Jithin Vijayan

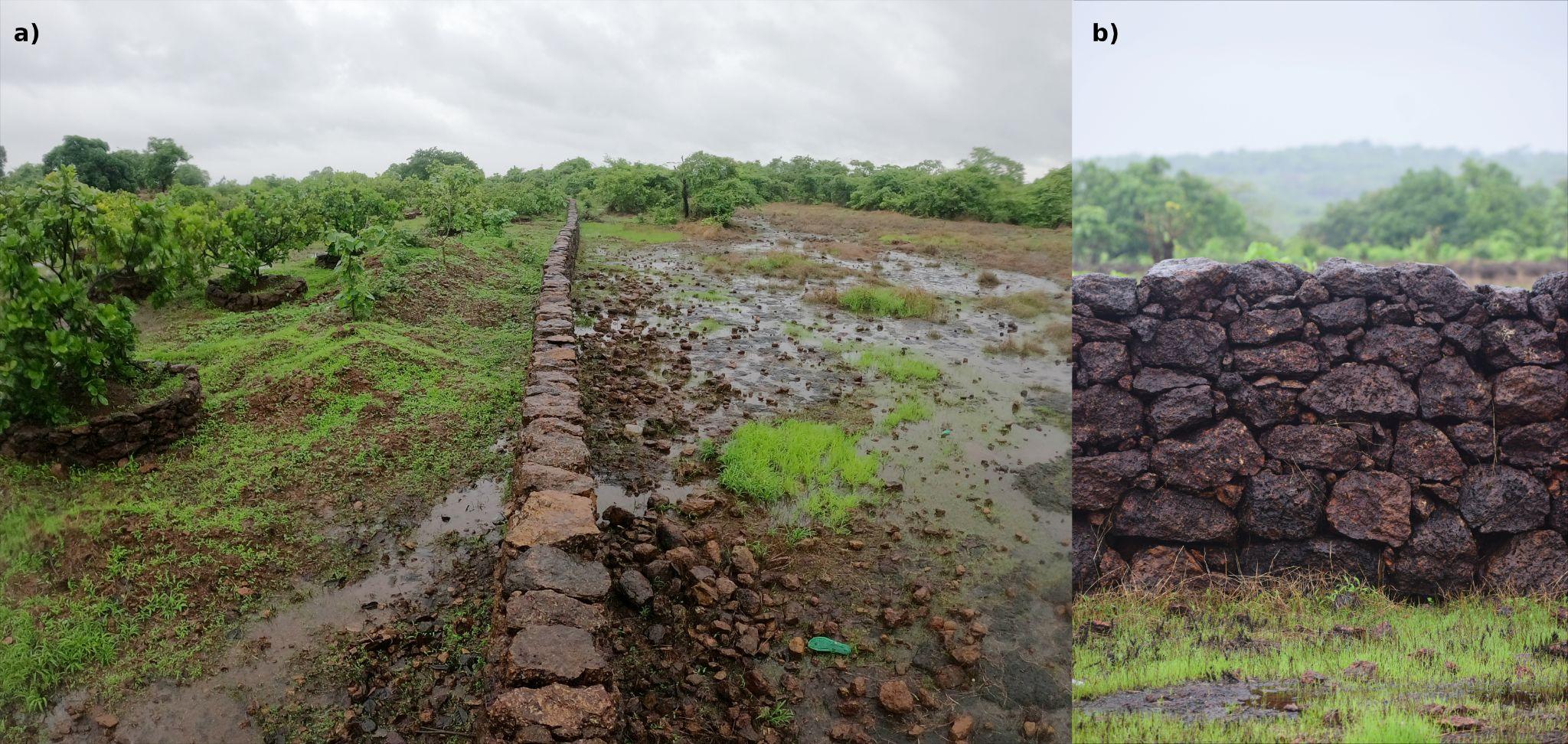

**Figure S6.** Orchards with a) rock walls along the boundary with undisturbed plateau; b) closeup view of a rock wall. Photographs: Jithin Vijayan

**REFERENCES**

1. Cortes-Gomez, A. M., Ruiz-Agudelo, C. A., Valencia-Aguilar, A., & Ladle, R. J. (2015). Ecological functions of neotropical amphibians and reptiles: A review. *Universitas Scientiarum*, *20*(2), 229–245. https://doi.org/10.11144/Javeriana.SC20-2.efna
2. Dalmolin, D. A., Tozetti, A. M., & Pereira, M. J. R. (2020). Turnover or intraspecific trait variation: Explaining functional variability in a neotropical anuran metacommunity. *Aquatic Sciences*, *82*(3), 62. https://doi.org/10.1007/s00027-020-00736-w
3. Dehling, J. M., & Dehling, D. M. (2021). Conserving ecological functions of frog communities in Borneo requires diverse forest landscapes. *Global Ecology and Conservation*, *26*, e01481. https://doi.org/10.1016/j.gecco.2021.e01481
4. Garg, S., & Biju, S. D. (2017). Description of four new species of Burrowing Frogs in the Fejervarya rufescens complex (Dicroglossidae) with notes on morphological affinities of Fejervarya species in the Western Ghats. *Zootaxa*, *4277*(4), 451–490. https://doi.org/10.11646/zootaxa.4277.4.1
5. Hernández-Ordóñez, O., Santos, B. A., Pyron, R. A., Arroyo-Rodríguez, V., Urbina-Cardona, J. N., Martínez-Ramos, M., Parra-Olea, G., & Reynoso, V. H. (2019). Species sorting and mass effect along forest succession: Evidence from taxonomic, functional, and phylogenetic diversity of amphibian communities. *Ecology and Evolution*, *9*(9), 5206–5218. https://doi.org/10.1002/ece3.5110
6. Olalla-Tárraga, M. Á., Diniz-Filho, J. A. F., Bastos, R. P., & Rodríguez, M. Á. (2009). Geographic body size gradients in tropical regions: Water deficit and anuran body size in the Brazilian Cerrado. *Ecography*, *32*(4), 581–590. https://doi.org/10.1111/j.1600-0587.2008.05632.x
7. Paluh, D. J., Stanley, E. L., & Blackburn, D. C. (2020). Evolution of hyperossification expands skull diversity in frogs. *Proceedings of the National Academy of Sciences*, *117*(15), 8554–8562. https://doi.org/10.1073/pnas.2000872117
8. Pittman, S. E., Jendrek, A. L., Price, S. J., & Dorcas, M. E. (2008). Habitat Selection and Site Fidelity of Cope’s Gray Treefrog (Hyla chrysoscelis) at the Aquatic-Terrestrial Ecotone. *Journal of Herpetology*, *42*(2), 378–385. https://doi.org/10.1670/07-1702.1
